# A testis-specific Heme Peroxidase HPX12 regulates male fertility in the mosquito *Anopheles stephensi*

**DOI:** 10.1101/2020.06.30.181388

**Authors:** Seena Kumari, Charu Chauhan, Jyoti Rani, Tanwee Das De, Sanjay Tevatiya, Punita Sharma, Kailash C Pandey, Veena Pande, Rajnikant Dixit

## Abstract

In humans, dysregulation of the antioxidant defense system has a detrimental impact on male fertility and reproductive physiology. Establishing such a correlation in disease vectors may pave a new way to manipulate male’s reproductive physiology, but remains the least attended. Since long - term storage of healthy and viable sperm earmarks male’s reproductive competency, we tested whether the anti-oxidative system protein also influences male fertility in the mosquito *An. stephensi*. We showed that a testis-specific HPX12 is critical for maintaining the physiological homeostasis of male mosquito’s reproductive organs. Disruption of this antioxidant enzyme by dsRNA silencing in the male mosquito severely impairs the reproductive potential of mated blood-fed female mosquitoes,resulting in a loss of 60% eggs. Our data demonstrate that increased ROS in the HPX12 mRNA depleted mosquitoes is an ultimate cause of sperm disabilities both qualitatively as well as quantitatively.

## Introduction

Mosquitoes are medically important insect pests because they transmit various infectious diseases such as malaria, dengue, chikungunya, andZika virus, etc. Seasonal variation has a direct impact on the mosquito’s population abundance and hence the disease transmission. Although the use of chemical insecticides still holds the key to the success of the vector control program, growing resistance urges the design of new alternative tools(1).Sincethe high reproductive capacity of these mosquitoes contributes to their role as disease vectors, strategy to disrupting sex-specific reproductive physiology could be an attractive tool to combat vector-borne diseases (2–5).

Mosquitoes are evolved witha more complex nature of mating behavioral properties and reproduction(6).During mating, both male and female mosquitoes undergo guided behavioral eventssuch as air swarming, effective coupling and correct reproductive organ positioning, rapid sexual engagement, and insemination. However, post-emergence, to outcompete male mosquitoes need to gain puberty, maintain healthy matured sperm and show behavioral competency in selecting, engaging and copulating in the crowd during swarming(7–9). Post-insemination,the male’s transferred seminal fluid proteins serve as powerful modulators of many aspects of female physiology and behavior, including re-mating, longevity, and egg production(10–14). Thus,post-mating adult females need to maintain long-term storage and survival of healthy sperms in their spermathecaforgonotrophic cycles(15).Maintaining healthy and viable sperms in unmated adult malesand/or post-mated females is crucial to the mosquito’s reproductive success, unraveling the basic mechanism could be targeted to reduce the fertility of the field mosquito population.

In vertebrates, including humans, studies have showna strong correlation between oxidative stress and infertility causes in males(16–19).In sperm cells,a substantial number of PUFA (polyunsaturated fatty acids) in their membrane and the epithelial presence of the Duox family servesas a major source of ROS (reactive oxygen species) generation in both vertebrates and invertebrates (20). While small amounts of ROS are required for normal sperm functioning, disproportionate levels can negatively impact the quality of spermatozoa and impair their overall fertilizing capacity (21). Uncontrolled production of ROS impairs the antioxidant system, causingincreased oxidative stress, which affects spermatozoa activity, damages DNA structure, and accelerates apoptosis, sperm motility, viability, and numbers(22–24). Thus, the evolution of a strong anti-oxidative system comprising enzymes such as heme peroxidase (HPX), Duox, catalase, SOD, GST, and other non-enzymatic factors, is believed to play a key role in maintaining optimal homeostasis for long-term storage and survival of sperms (25,26). But the mechanism of this regulation remains unknown in mosquitoes until a recent demonstration that HPX15 activation ensures not only survival of *Plasmodium* in the gut(27)alsohealthy sperm in the spermathecal of mated adult female *An. gambiae* (28). Since males do not feed on blood, aiming either at a reduction of mosquito populations or population replacement, male-specific transcripts/proteins may serve as potential targets.

*Anopheles stephensi* is a major malarial vector that transmits at least 20% of urban malaria in India. The increased challenge of using IRS/LLINs due to insecticide resistance, larval control is recommended in urban settings (29), but it does not suffice to control adult mosquitoes. Recently, we reported that mRNA depletion of a salivary specific hemeperoxidase HPX12 (*AsHPX12*) altershost-seeking abilities of adult female mosquitoes (30). Now, here we showthat how the testis-specific*AsHPX12* is essential to maintain male’s fertility and reproductive physiology in this mosquito species.In a combination of transcriptional profiling and functional analysis, we demonstrated that disruption of HPX12 by dsRNA silencing also reducesthe egg-laying capacity of the adult female mosquitoes. Our results further highlight that sperm function can be impaired to disrupt mosquito vector reproductive physiology, affording opportunities for vectorcontrol.

## Material and methods

Fig. S1 (supplemental file) represents an overview of the technical design and experimental workflow.

### Basic Entomology

#### Mosquito rearing

*An. stephensi* mosquitoes were reared in cage under standard conditions (26°C--28°C, 65%-80% relative humidity, 12:12h light) (31,32). Eggs were floated in a pan filled with deionized water and once larvae hatched, approximately 1000 larvae were reared in an iron tray (66cm × 45cm × 17cm). Larvae were fed on a 1:1 mixture of dog food (Pet Lover’s crunch milk biscuit, India) and fish food (Gold Tokyo, India) as before. Post-emergence adult mosquitoeswere fed daily on sterile sugar solution (10%) using a cotton swab, and for routine Oviposition and gonotrophic cycle maintenance, the rabbit was offered a blood meal. All protocols for rearing and maintenance of the mosquito culture were approved by the ethical committee of the institute.

#### Pupae sexing

Adult mosquitoes were separated by sex as pupae using a light microscope and placed in separate cages in dishes filled with deionized water to maintain virginity. Cages were inspected when adults emerged and any of the wrong sexed adults were removed.

#### Mosquito mating

Mating assays were performed to determine the percentage of females mating throughout one night. For 100 virgin females, 3-4 days after eclosion, were kept in a cage containing over 100 virgin and age-matched male mosquitos and put overnight in the dark. Subsequently, we used a single overnight mating method in all experiments. Mating success was assessed by dissecting and visualizing live sperm in spermathecae of randomly selected 8-10 mosquitos, in addition to the verification through sperm-specific primers (ams, mts) by RT-PCR in Vir vs. mated spermatheca.

### Molecular Biology

#### RNA extraction, cDNA preparation, and quantitative RT-PCR

Experimentally required tissues were dissected and pooled from the cold anesthetized adult female and male mosquitoes under different conditions i.e. virgin, mated, was injected. To examine the tissue-specific expression of target genes, selected tissues such as hemocyte, spermatheca, salivary gland, male reproductive organ (male accessory gland, testis) were dissected from age-matched ‘control’ vs. ‘test’ mosquitoes. Total RNA from collected tissues was isolated using the standard Trizol method as described previously (31).RNA was quantified by using a NanoDrop Spectrophotometer (Thermo Scientific). Isolated ~1μg total RNA was utilized for the synthesis of the first-strand cDNA using a mixture of oligo-dT and random hexamer primers and Superscript II reverse transcriptase as per the described protocol (Verso cDNA synthesis Kit, Cat#AB-1453/A, EU, Lithuania; Sharmaet al., 2015a,b). For differential gene expression analysis, routine RT-PCR and agarose gel electrophoresis protocols were used.

The relative abundance was assessed using SYBR Green qPCR master mix (Thermo Scientific), using a CFX96 PCR machine (Bio-Rad, USA). PCR cycle parameters involved an initial denaturation at 95°C for 15 min, 40 cycles of 10 s at 95°C, 15 s at 52°C, and 22s at 72°C. After the final extension step,melting curves were derived. Each experiment was performed in three independent biological replicates. The relative quantification results were normalized with an internal control (Actin), analyzed by the 2–^ΔΔ^Ct method.

#### Gene silencing RNAi assays in adult mosquitoes

For the knocking down of *AsHPX12*, dsRNA primers carrying T7 overhang were synthesized as listed in the supplemental table (ST2). The amplified PCR product was examined by agarose gel electrophoresis, purified (Thermo Scientific Gene JET PCR Purification Kit #K0701), quantified, and subjected to double-stranded RNA synthesis using Transcript Aid T7 high-yield transcription kit (Cat# K044, Ambion, USA),andthe dsrLacZ gene was used as a control. Approximately ~69 nl (3 μg/μl) of purified *dsRNA* product was injected into the thorax of a cold anesthetized 1–2-day old male mosquito using a nano-injector (Drummond Scientific, CA, USA). The silencing of the respective gene was confirmed by quantitative RT-PCR after 3-4-days of dsRNA injection.

### Microscopic assay

#### Male reproductive organ (MRO) morphological examination

After participating in the experiments, live males were cold anesthetized, placed in a drop of phosphate-buffered saline (PBS) over a microscopic glass slide, and examined under a dissecting microscope (16×). Their reproductive system was removed using needles to pull out the last segment of their abdomen. Slow excision removed the whole male reproductive system, including the testes, accessory glands, and ejaculatory duct. Subsequent observations were made under a compound microscope (40×) to visualize the ultrastructure of male reproductive organs, including the testes and accessory glands. The images of the target tissue/organ were captured with the assistance of a Smart-Phone camera (with zoom option/12X), stabilized on the eyepiece of the microscope. The final images of both the control and test groups, captured in identical magnifying parameters, were further processed using professional software (Microsoft Photo Edit software;version 2020.19111.24110.0). Likewise, the motility of the sperms and their associated fluid movement was recorded under the camera/video option, and processed for trimming and clipping through professional video editing software.

#### Trypan blue staining

The altered morphology of the spermatozoa(testis and seminal vesicles)was examined by trypan blue, which could differentially stain dead or alive cells (33). To do this, the testis and seminal vesicle from dissected MRO was carefully separated with a microneedle and collected in a 1.5 ml tube containing 100μl PBS. After manual homogenization with a gentle pressure enough to rupture the testis. A drop of ~20 μl sample of spermatozoa containing seminal fluid was mixed with 10 μl h Trypan blue solution (0.27%) on a slide and examined microscopically.

#### ROS Determination assay

To determine the level of ROS generation, MRO (MAG, Testis and seminal vesicles) from HPX12 silenced or control mosquitoes were collected in PBS, and continue to incubated with a 2 mM solution of the oxidant-sensitive fluorophores, CM-H2DCFDA [5-(and-6)-chloromethyl-29,79-dichloro-dihydrofluorescein diacetate, acetyl ester] (Sigma) on a microscopic glass slide. After a 20-min incubation at room temperature in the dark, the MRO was washed three times with PBS. Next, the MRO was transferred to a new glass slide in a drop of PBS and checked the fluorescence intensity under a fluorescent microscope at 490 nm (FITC filter).

#### Fertility Assays

Overnight mated females (mated either with control male, or HPX12 silenced male mosquitoes) were blood-fed over a rabbit. After bloodfeeding, partial/incomplete blood-fed females were removed, while restfully engorged females were kept in a cage for 48hrs, and then individually placed in netted plastic cups containing half-filled water and fixed with blotting paper on which to lay eggs (day 4). After 24hrs. The total number of eggs and the total number of hatched eggs (broken eggshell) were counted, and the proportion of infertile eggs was calculated.

#### Statistical analysis

Statistical analysis was performed using Origin8.1. All these data were expressed as mean ± SD.Differences between test samples and their respective controls were evaluated by the paired Student’s *t*-test for transcriptional profiling.The*Mann –Whitney U test* was used to evaluate and analyze egg-laying experiments, where results were considered significant if the *p*-value was less than 0.05.Each experiment was performed at least thrice to validate the findings.

## Results

### Mating alters HPX12 gene expression in male reproductive organs

Because of a very high rate of cell division and mitochondrial oxygen consumption in testicular tissue, the onset of the spermatogenesis process is highly susceptible to oxidative stress during the early development of male gonads (29). Therefore, first, we evaluated the expression pattern of at least 11 transcripts encoding the antioxidant system (AOS) associated with HPX as well as other family proteins during the aquatic development of the mosquito.All tested genes showed significant modulation throughout development but were exceptionally enrichedin the pupae compared to other stages (Fig.1a; also see SupplementalFig.2a).Similarly, a higher expression of AOS genes was observed in newly emerged males than in female mosquitoes (Supplemental Fig.2a), where HPX12 further showed >5-fold (p<0.00045) higher expression in the male reproductive organ (MRO) than midgut, salivary glands and hemocytes (Fig.1b). Next, though, we observed an unusual pattern coinciding with enriched expression of all HPX members during young (3-5Days) or aged (12D) mosquitoes(Fig. 1c); however, surprisingly,the expression of *AsHPX12* exceptionally remains higher than other tested AOS genes in unmated virgin male MRO (Supplementary Fig.2b,c). Finally, after mating a significant loss in the expression level/(p<0.0003) (Fig.1d), correlate HPX12 may likely have a potential role in oxidative stress tolerance of the male mosquito’s reproductive system.

**Fig.1.**
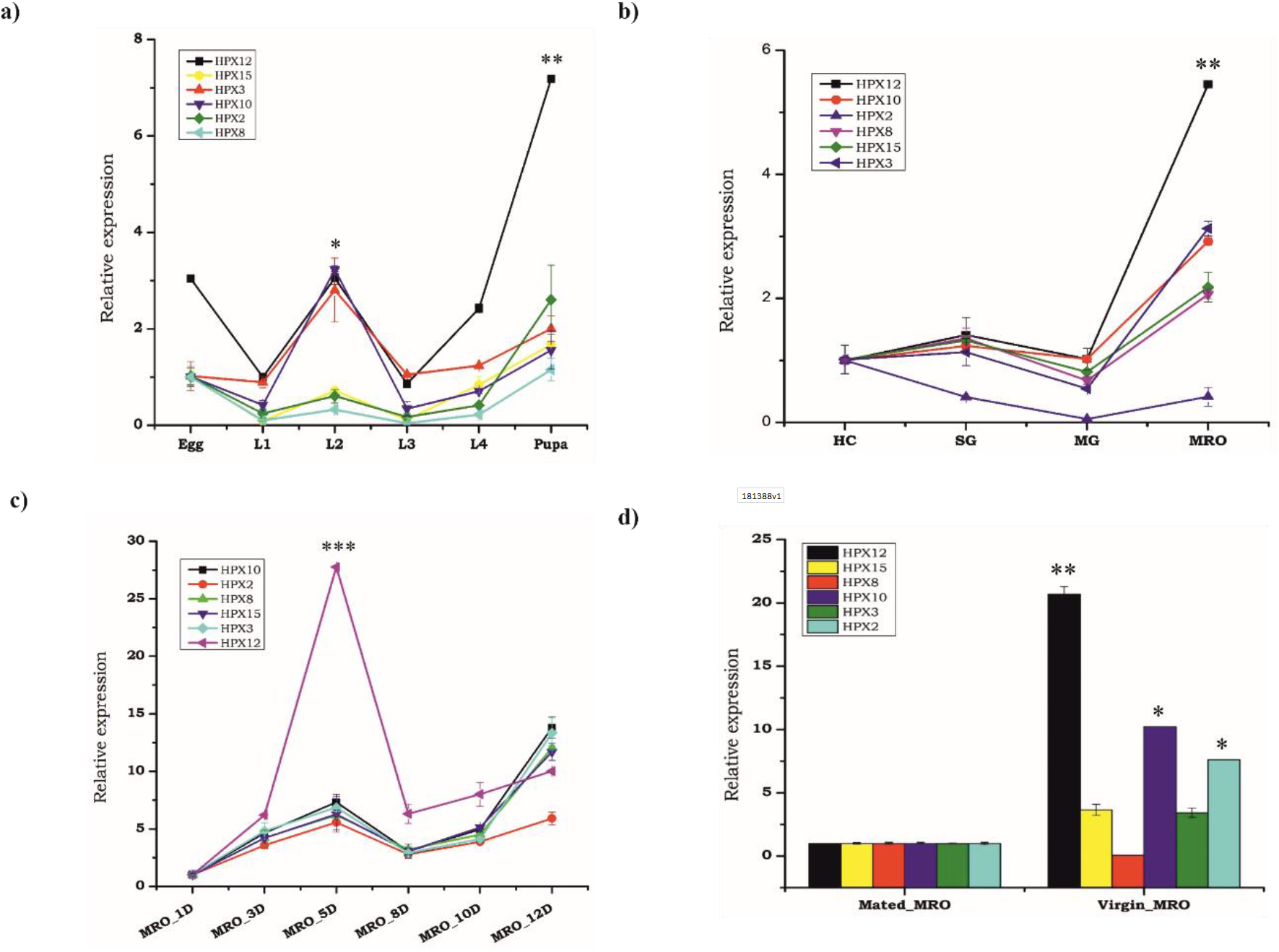
Transcriptional profiling of the heme peroxidase family protein transcripts (hpx) in mosquito *An. stephensi* **(a)** Heme peroxidases (HPX12,10,15,3,8 and 2) expression during embryonic development: L1-larval instar one, L2-larval instar 2, L3-larval instar 3 and L4-larval instar forth, pupa (p<0.0001) (n= 3,N10);**(b)** Tissues specific expression kinetics of HPX family in male mosquito tissues SG: Salivary glands; MG: Midgut; HC: Hemocytes, MRO; male reproductive organs (p<0.00045); **(c)** Age-dependent expression of HPX family in virgin male mosquito reproductive organ (MRO) i.e. 1D(day),3D,5D (p<0.0008564),8D and 10D,12 days; **(d)** Mating-induced changes in hpx family in male reproductive organ Vir vs. Mated (p<0.00324). Three independent biological replicates (n=30, N3) were considered for statistical analysis viz: *p<0.05; **p<0.005 and ***p<0.0005 using Student’s *t-test*.(*n*=represents the number of mosquito pooled for sample collection; *N*= number of replicates)

### HPX12 influences sperm hosting testis physiology

To test and establish whether HPX12 influencessperm physiology, initially,we examined the co-expression pattern of sperm-specific markers, namely AMS/MTS, in the testis of aging adult male mosquitoes (Supplemental Fig.3a).Surprisingly, perfect alignment with HPX12 expression pattern in both mated and unmated mosquito (SupplementaryFig3b), and aquantitative loss in the sperm number, as measured by AMS/MTS expression in the HPX12 mRNA depleted mosquitoes (p<0.008774), suggesting a key role of HPX12 in testicular homeostasis maintenance(Fig. 2b). When mated with HPX12 silenced male mosquitoes, adult female mosquitoes showed a 60% reduction in their egg number (Fig. 2c,d), confers indicating that *HPX12*is key to regulating the male mosquito’s sperm fertility and testis physiology.

**Fig. 2:**
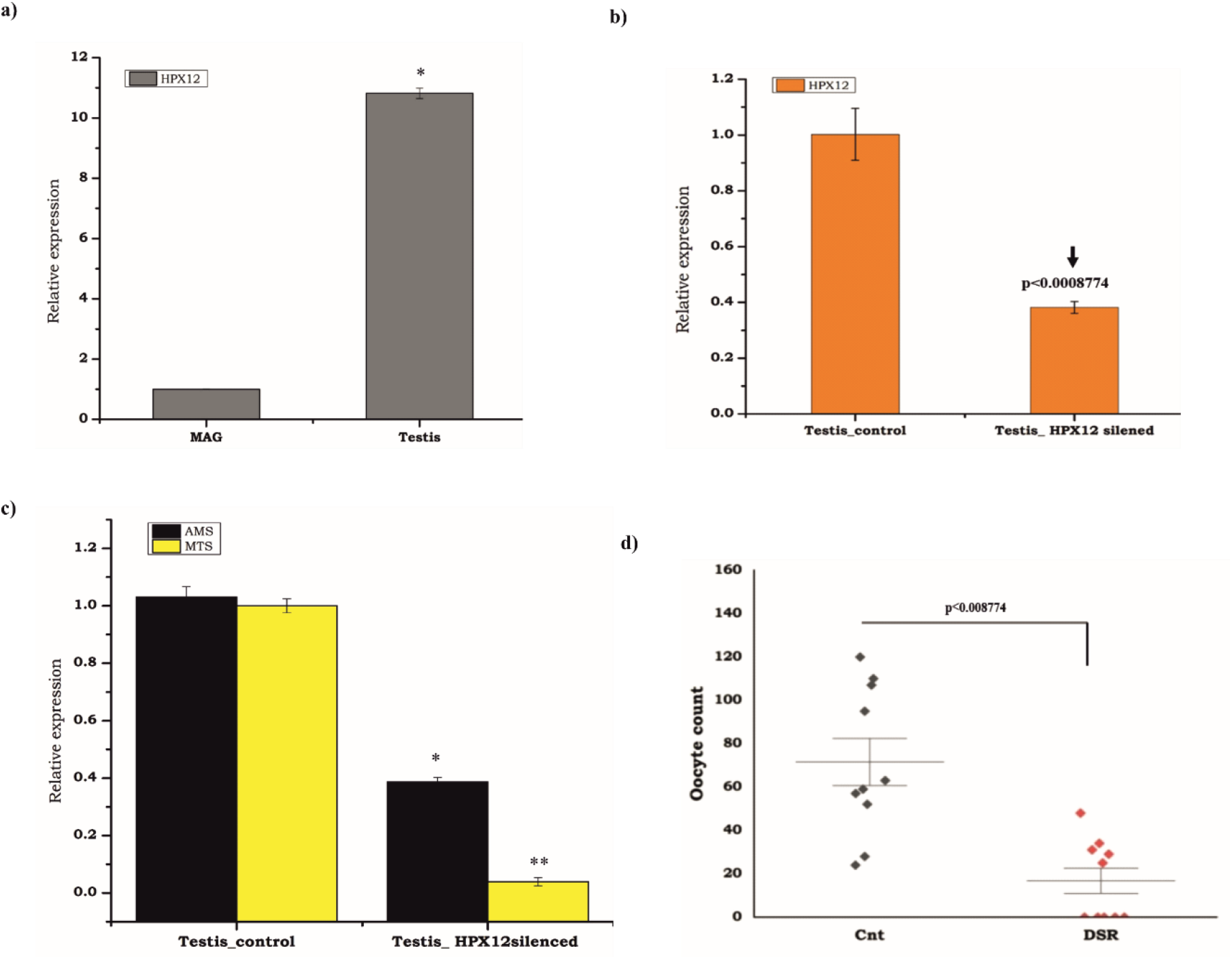
HPX12 abundantly express in the testis and required to prevent infertility: (a) Hpx12 expression in male mosquito reproductive organs(MAG: male accessory gland and testis (p<0.01504); (b) HPX12 silencing exhibited a >70% reduction in mRNA level as compared to control mosquito (p<0.0008774); (c)Co-expression of sperm-specific genes (ams, mts) in the control and HPX12 silenced mosquitoes; (d) Mating with HPX12 silenced male mosquitoes reduces the egg-laying capacity of adult female mosquitoes:-the percentage of eggs that are infertile laid by females injected with ds*LacZ.* Females were allowed to lay eggs between days 4 and 5. (median with range; N=3-5, n=15–25 females/trial) and (p<0.008774, Mann-Whitney *U* test). A minimum of three biological replicates were performed.

### HPX12 regulates sperm fertility

Oxidative stress resulting in the production of reactive oxygen species (ROS), may have both positive and negative impacts on sperm functions, such as motility, viability, and loss of fertility(33,34). Previously, the heme peroxidase HPX15 enzyme has been shown to help in long-term storage and survival of sperms in the female mosquito’s spermatheca (28). Since HPX12 silencing in male mosquito reduced female reproductive outcome, we hypothesized that it may be due to altered quality/defective sperms, and therefore tested it in male mosquito’s testes.Our initial phase-contrast microscopic and video capture analysis evidenced a significant loss in viability and mortality of the sperm in the HPX12 silencedthan control male mosquito’s group (Fig.3a,b,c/VideoS1,S2).Compared to the control, a significant loss in the integrity of the testis was readily evident while dissecting and removing itfrom HPX12 silenced mosquitoes (Fig.3a). A parallel increase of dead sperm, visualized through trypan blue, which marks the dead cell (sperm head), possibly due to loss in the motility or distortion/abnormality (aberrant sperm) of the tail (Fig.3b,c).

**Fig.3.**
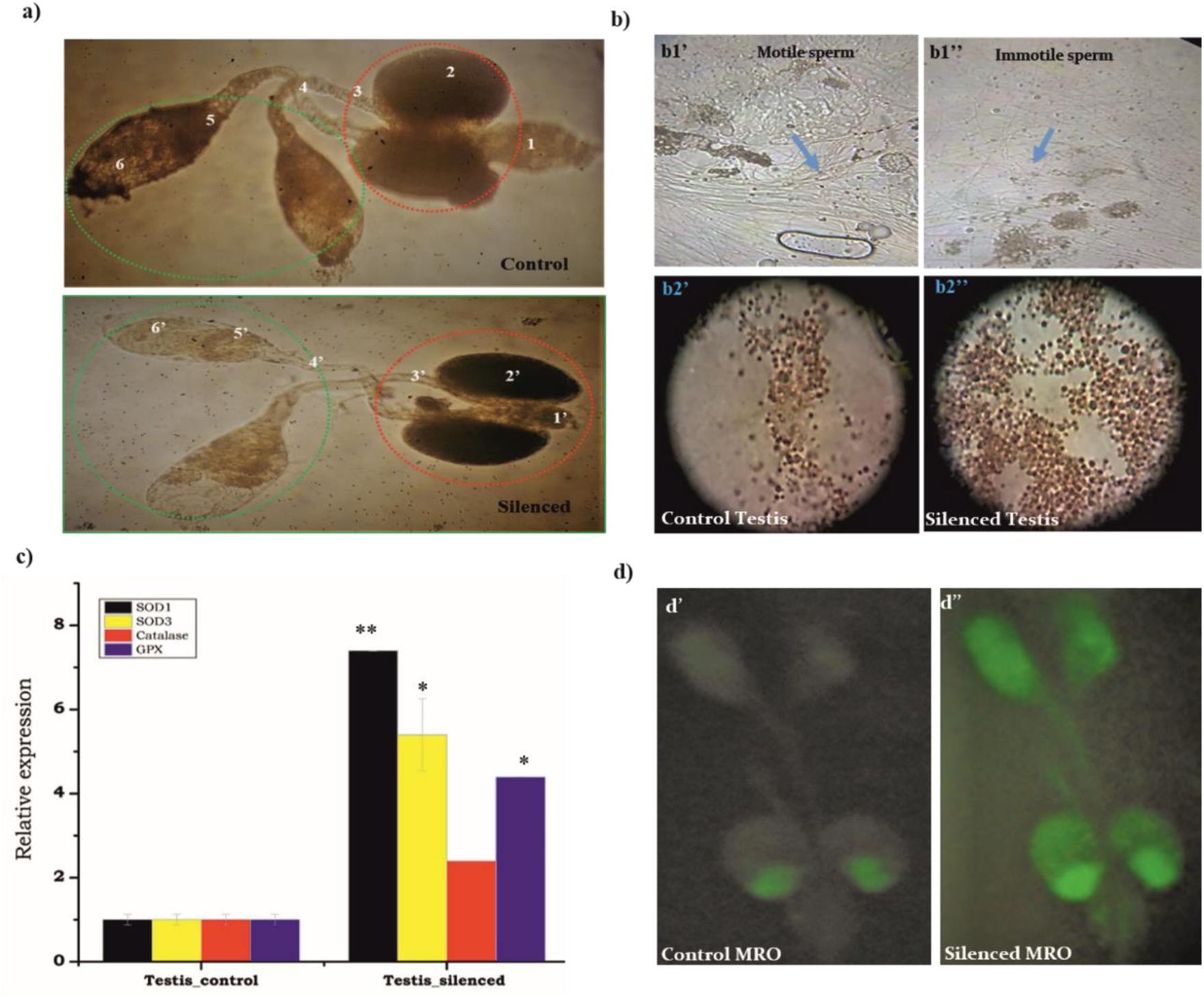
HPX12 silencing affects sperm physiology (morphological and molecular) in male reproductive organs (mag+testis); **a’)** The reproductive system of control and silenced male mosquitowere dissected and placed in a drop of phosphate-buffered saline (PBS) andexamined under a dissecting microscope (16×). The male reproductive organ features i.e. numbers on the figure represented as a’ 1 -ejaculatory duct, 2-accessory glands, 3-seminal vesicle, 4-vas efferentia 5 - sperm reservoir, and 6 – spermatocytes; **(a’’)** Once HPX12 knocked down, they were placed in a drop of phosphate-buffered saline (PBS) and examined under a dissecting microscope (16×). The reproductive system of male *An. stephensi* showing features i.e. numbers on the figure represent; a’’ 1’-ejaculatory duct, 2’-accessory glands, 3’-seminal vesicle, 4’-vas efferentia 5’-sperm reservoir, and 6’ – spermatocysts (reduce size). **(b)** Effect of HPX12 silencing on sperm quality(motility), examined under a light microscope (40×), male testis/seminal vesicles from control (b1’) and HPX12 silenced (b1”) mosquito group,were visualized in PBS, highlighting motile (curved shaped) healthy and abnormal sperm structure (light blue arrow), respectively (also see videosS1, S2); (**b2’-b2”)** Tryptan blue stained testis of male mosquito showingincreased dead/abnormal structure of sperm bundles in the silenced mosquito group (**b2”)** than control group (**b2’)**; **(c)** Relative antioxidant enzyme transcripts expression in response to increased oxidative stress after HPX12silencing:-SOD1 (p<0.00072),SOD3(p<0.042),catalase(p<0.587) and GPX (p<0.0085);**(d)** Effect of HPX12 silencing on ROS generation in the male reproductive organ:-Control/**d’** (ds*LacZ*), and ds*Hpx12/***d** “injected virgin male mosquito’s MRO were incubated with DCFDA, a dye leading increased fluorescence intensity in response to ROS.

The up-regulation of all selected ROS-generating transcripts, namely SOD1 (p<0.0097), SOD3(p<0.042), and catalase(p<0.587),and increased fluorescent intensity in the MRO, as determined by DCFDA assay, in the HPX12 silenced mosquito group ensured thataltered ROS levelshavea detrimental impact on the loss of sperm quality (Fig.3a/b). Together,these data confirmed that HPX12 mRNA depletion may severely impair the anti-oxidative defense system and cause infertility responses by altering the quantitative and qualitative nature of sperms.

### HPX12 also influences Accessory Gland Proteins (ACPs) /TEP1 gene expression

Male seminal fluid proteins containing accessory proteins (ACPs) are key modulators of many aspects of female reproductive and behavioral physiology, such as re-mating, longevity, and egg production. Male’s successful copulatory event of ejaculation is a result of an optimal mixing of seminal fluid from the accessory gland (MAG) and sperms from the testis (Supplementary Fig.4a,b). However, unused sperms in aging mosquitoes are removed by the action of phagocytosis activity of immune factors such as TEP1(4). Therefore, we tested whether HPX12 has any functional correlation with ACPs and TEP1 gene expression. Surprisingly, we observed that both mating or HPX12 mRNA silencing caused a significant down-regulation of tested ACPs as well as TEP1 expression, suggesting that HPX12 also has an indirect influence on ACPs/TEP1 to preserve healthy sperm and high fertility rates in the male mosquitoes (Fig. 4a, b).

**Figure 4.**
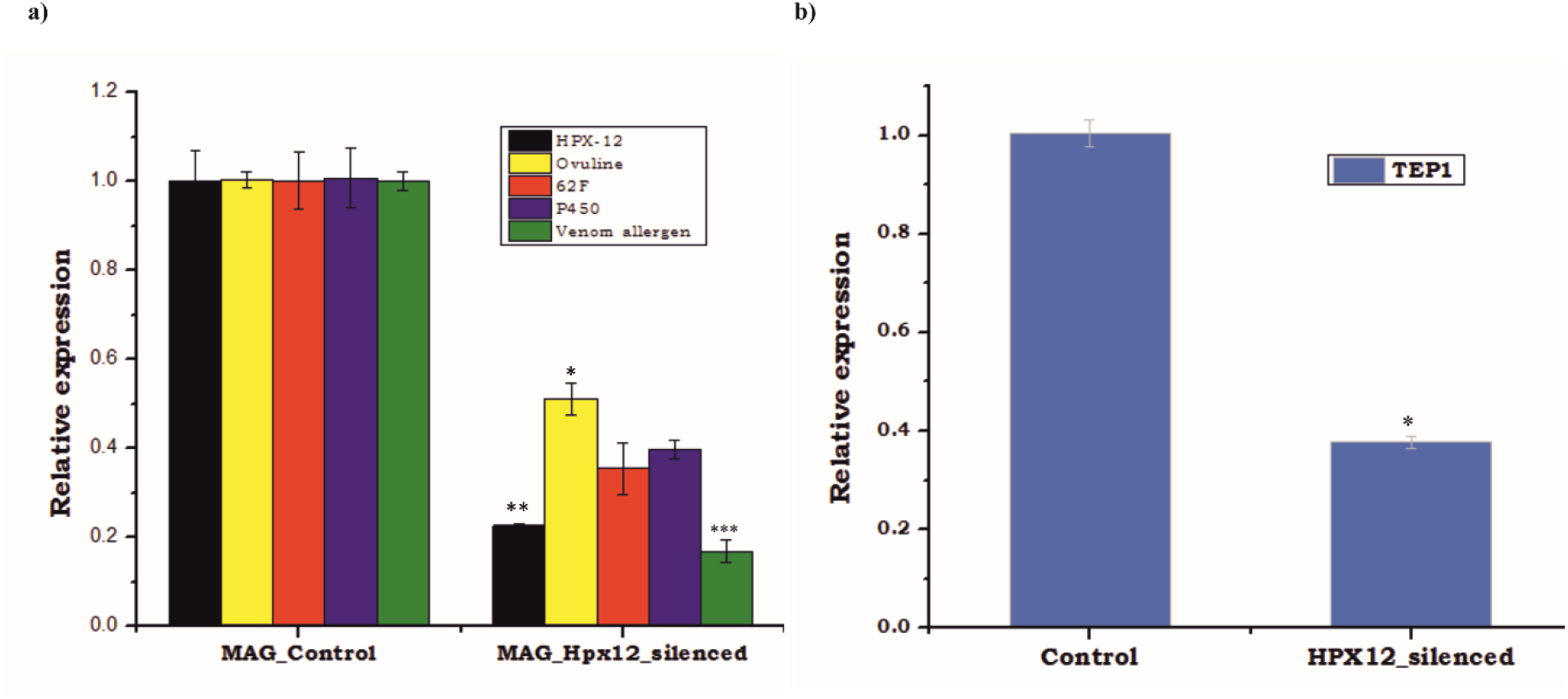
Relative levels of accessory gland proteins and TEP1in Hpx12 silenced male MRO: depletion on male fertility a) After silencing, accessory gland proteins Ovuline (p<0.0148),62f (0.033068), P450(p<0.0188012) andvenomallergen (p<0.000375),) depletions decreased sperm storage and protection. b)The proportion of testes with apoptotic cells was examined for TEP1 (p<0.004345)transcripts in HPX12 silenced and control male mosquito, N3,n=30. (*n*=represents the number of mosquito pooled for sample collection; *N*= number of replicates)

## Discussion

Our findings highlightthe transcriptional modulation of anti-oxidative stress-responsive proteins during mating, and the crucial role of *AsHPX12* in maintaining male mosquito’s fertility. Through functional analysis, we identified*AsHPX12*as a possible target to reduce male mosquito’s fertility in field populations. Due to high susceptibility to reactive oxygen species (ROS), sperm production and maintenance in the gonad is critical to the male’s reproductive system (21,35–37).A positive correlation between levels of reactive oxygen species (ROS) and sperm fertilityis welldocumented in invertebrates(20,38,39),but lack in mosquitoes. Previous studies have suggested that the maturation of sperm mother cells i.e. spermatids, coincides with the early development of the gonad system of male mosquitoes(40).

Our observation of enriched anti-oxidant system geneexpressionfurther supportsthe idea that gonads may mature during the earlypupal stage(41,42). However, newly emerged male mosquitoes mayremain unfit for sexual courtship until it achievesa morphological change in the external genitalia through inversion of the terminalia within the first 24 hrof emergence(43). In many Anopheline mosquito species, male accessory glands mature during the first few days and usually attain optimal puberty at the age of 5–7-daysof adult life (7,44).Anage-dependent AOS gene expressiondata analysis and their alteredexpression after mating, together confers that laboratory-reared adult male *An. stephensi* may achieve puberty age within 3-5day after emergence, where an active anti-oxidative defense system is necessary to maintain healthy sperm until mating event completion.

Many studies in humans and other vertebrates strongly suggest that dysregulation of the antioxidant system and imbalance of reactive oxygen species (ROS) productionis the main cause of male infertility (19). Although such a relationship has not been studied in any male mosquito species, HPX15 has been found crucial for long-term storage and survival of sperm in the spermathecaof mated adult female mosquito *Anopheles gambiae*(28). Exceptionally elevated levels of HPX12 expression prompted testing its potential role in male mosquito’s reproductive physiology and fertility maintenance in *An. stephensi*. We noticed that effective silencing of HPX12 (>80% reduced mRNA level)not only caused significant loss of sperm but also reducedthe egg-laying capacity of the gravid adult female mosquitoes.Failure to combat the increased ROS in the HPX12 silenced mosquito’s testis, and parallel loss of sperm viability and motility, validate the hypothesis that testis-specific HPX12 is key to prevent the damage of sperms in adult male mosquitoes. Furthermore, altered expression of accessory proteins (ACPs), which contribute a major content of seminal fluids, and TEP1 protein, which helps in the removal of damaged sperms, together may also affect the male mosquito’s reproductive competency(4). We correlate HPX12 could protect sperm cells directly, similar to mating induced expression of heme-peroxidase HPX15 into the spermathecal lumen of the mated female mosquito *An. gambiae*(28).

Previously, we characterized HPX12 as a salivary specific heme peroxidase family member HPX12, originally identified from *Plasmodium vivax-*infected salivary RNAseq data of the adult female mosquito *Anopheles stephensi*. We showed that salivary HPX12 silencing severely impairs pre-blood meal-associated behavioral properties such as probing time and probing propensity, possibly by altering physiological-homeostasis of the salivary gland. Although the exact mechanism is yet to unravel, we proposethat HPX12 could serve as a suitable molecular target to manipulate the behavioral physiology of host-seeking in adult female mosquitoes, and the reproductive physiology of adult male mosquitoes to design asex-specific vector control strategy.

## Conclusion

Studies in humans strongly prove that a controlled regulation of the anti-oxidative defense system has a direct influence on male fertility and reproductive success. Here, we tested this hypothesis in mosquito *An. stephensi*, and demonstrated that dysregulation of thetestis-specific heme-peroxidase homolog *AsHPX12* severely impairs male fertility. Our data provide evidence that HPX12 is crucial for maintaining optimal homeostasis for storing and protecting healthy sperms in male reproductive organs.

## Supporting information

Control_Mosquito_Sperm_Motility

HPX12_Silenced_Mosquito_Sperm_Motility

## Acknowledgment

We would like to thank all the technical staff members of the central insectary for mosquito rearing and Kunwarjeet Singh for lab assistance. We are grateful to the malaria clinic facility support for their contribution *Plasmodium* infection study. Finally, we thank Xceleris Genomics, Ahmedabad for NGS sequencing.

## Grant Information

Work in the laboratory was supported by Indian Council of Medical Research (ICMR), Govt. of India.SK is the recipient of CSIR Research Fellowship (09/905(0015)/2015-EMR-1).

## Authors’ contribution

SK, RKD conceived scientific hypothesis and designed the experiments.CC, JR, ST, TDD, and PS Technical support for tissue dissection/collection and gene profiling, Data reviewing, and presentation.RKD, KCP contributed reagents/ materials/ Analysis tools; SK wrote the paper, KCP, RKD edit the MS. All authors read and approved the final manuscript.

## Conflict of Interest

No competing interests were disclosed.

## Notes

### Competing Interest Statement

The authors have declared no competing interest.

