## Supplemental data for "A testis-specific Heme Peroxidase HPX12 regulates male fertility in the mosquito *Anopheles stephensi*"

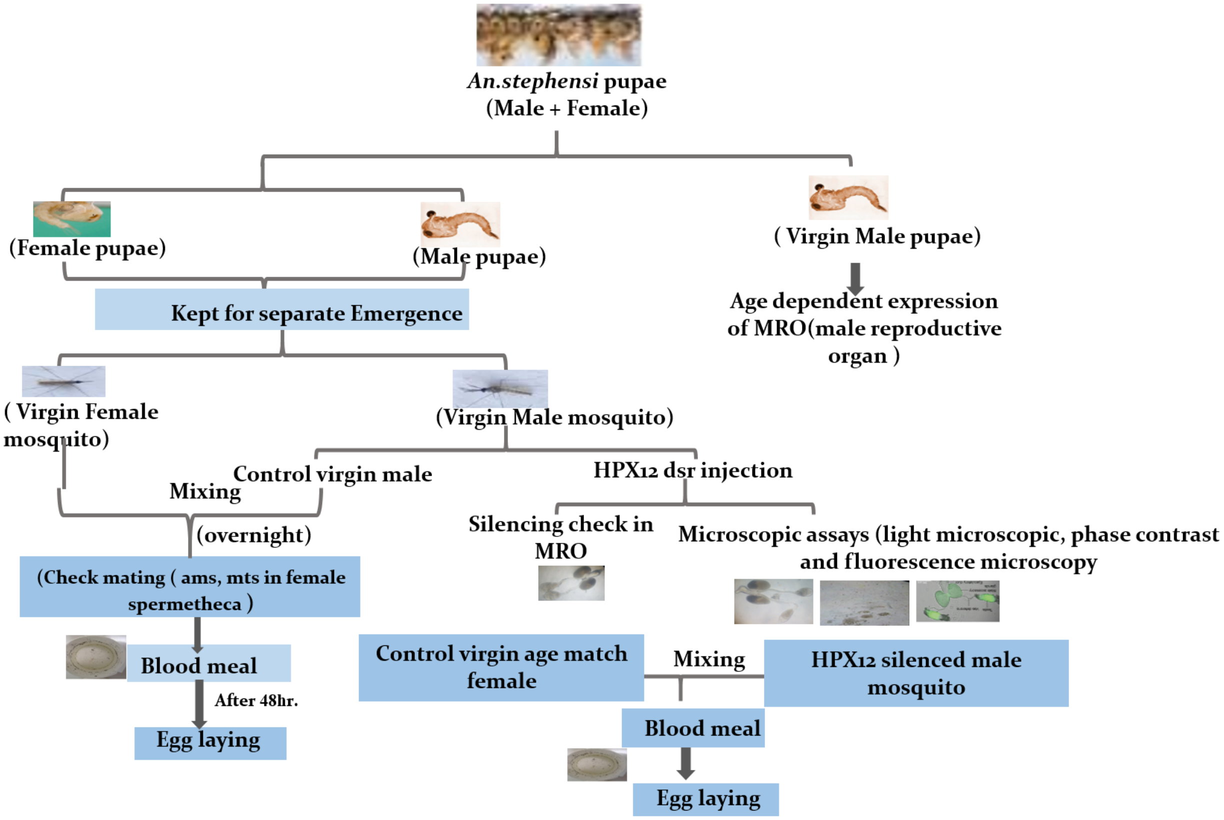


**Supplementary figure 1** Technical designing and experimental workflow

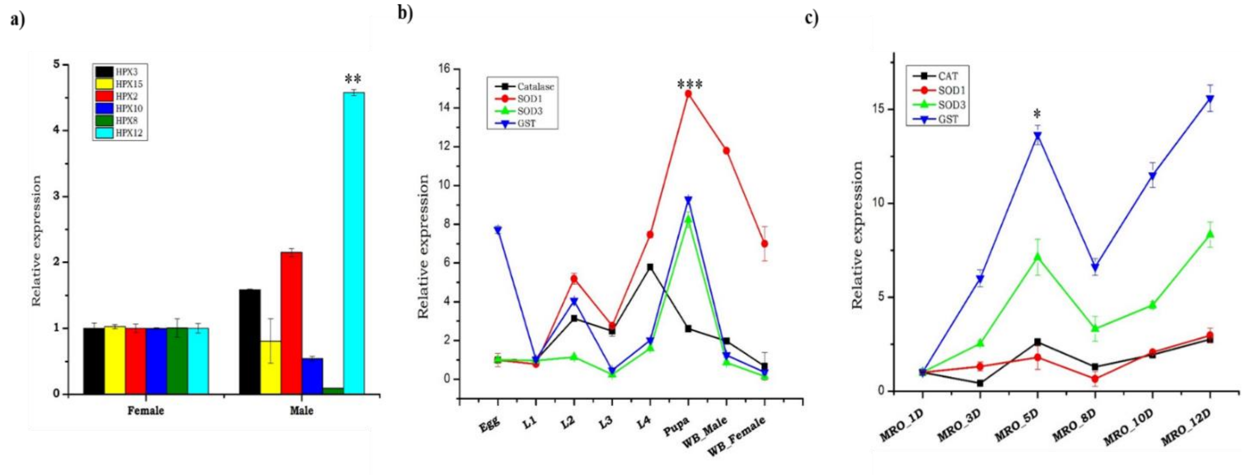


**Supplementry Figure.2** Transcriptional profiling of the AOS genes transcripts (hpx) in mosquito *An. stephensi* **a)** Heme peroxidases (Hpx 12,10,15,3,8 and 2 p<0.000388) expressions in male and female whole body (n=3, N1); b) AOS genes proliferation in aquatic stage i.e. L1- larval instar one, L2- larval instar 2, L3-larval instar 3 and L4-larval instar forth, pupa (p<0.0001, n= 3, N10); **c)** Age-dependent expression of AOS family in virgin male mosquito reproductive organ (MRO) i.e. 1D (day), 3D, 5D (p<0.008564), 8D and 10D, 12 days.

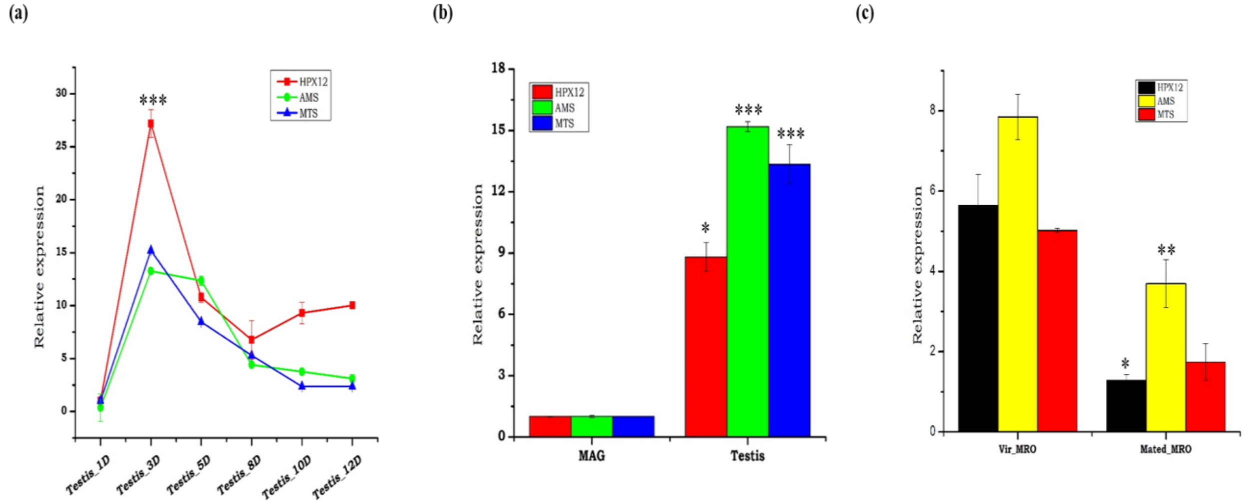


**Supplementary Figure.3** Transcriptional profiling of the HPX12 and sperm specific transcripts (AMS ,MTS) in mosquito *An. stephensi* **a)** Age-dependent CO -expression of hpx12 and ams, mts in virgin male testis , 1D ,3D,5D,8D, 10D and 12D. b) Transcriptional profiling of hpx12 (p<0.01504) and ams(p<0.000894), mts (5.57E-05) in male reproductive tissues (testis, mag) ; c) Mating-induced changes in hpx12 (p< 0.01010), mts (p<0.002276) and ams (p<0.035324) in male testis Vir vs. Mated. Three independent biological replicates (n=3, N30) were considered for statistical analysis viz. *p<0.05; **p<0.005 and ***p<0.0005 using Student's *t-test*.

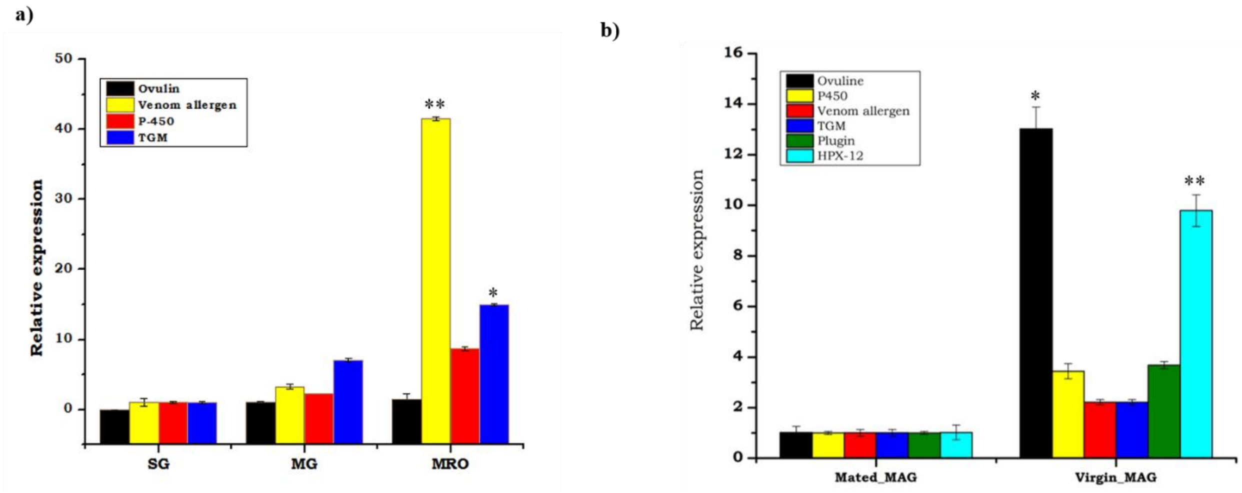


**Supplementary Figure 4** Transcriptional profiling of the Accessory gland proteins (ACPs) in mosquito *An. stephensi;* a) Tissue specific expression of Acps i.e. Ovuline, venom allergen ($p<0.000375$), P-450) TGM ($p<0.0190$); b) Mating-induced changes in hpx12 ($p< 0.01010$), Ovuline ($p<0.002276$) and P450 ($p<0.035324$), Plugin, venom allergen ,TGM in male MAG (male accessory gland ) Vir vs. Mated. Three independent biological replicates (n=3, N30) were considered for statistical analysis viz. *$p<0.05$; **$p<0.005$ and ***$p<0.0005$ using Student's *t-test*.

**Supplemententry table (ST1) Primer list :**

| S.No. | Gene Name | Primer Sequence |
|---|---|---|
| 1 | HPX2 | Fw: CACGAAGCTAAAAATTGTCC<br>Rev: AAGAGATGCTCCAGATCGTA |
| 2 | HPX3 | Fw: AGTTCTTCACGGTTCATCAC<br>Rev: GATCTTCTCCAGGTGTTTTG |
| 3 | HPX8 | Fw: AAGCTGTACCAAGAAGCTC<br>Rev: GCTCCTGATACTTCGTTAAA |
| 4 | HPX10 | Fw: AAGAAGGTTGACGGTACAGA<br>Rev: CAACGTTAGCAGTCGTATCA |
| 5 | HPX15 | Fw: CTATTGTCGCAACAGTACGA<br>Rev: CGGTAAACTGTCCATCATTT |
| 6 | Acp62f | Fw: 5' TACACGCTAGTGGTTTTGC 3'<br>Rev: : 5’ AGCAAGATGTGGAAGGAATA 3’ |
| 7 | HPX12 | Fw: GAACAGTGCCACCGATACCT<br>Rev: CCGAGATAATAGGGCAACCA |
| 8 | P-450 | Fw: TCATAATGTGGTGTACAAGC<br>Rev: ACCTAATCTCTATCGGTGTG |
| 9 | Catalase | Fw: 5' CCCAACTATTTCCCGAAC 3'<br>Rev: 5’CAGGTGTCCCACAATGTT3' |
| 10 | GST | Fw: 5' AACGGGTCGTCGATTACT 3’<br>Rev: 5' AGGTCGAACTGGAATGCT 3’ |
| 11 | DsrLacz | Fw: 5' TAATACGACTCACTATAGGGGAGTCAGTGAGCGAGGAAG 3' |

| | | |
|---|---|---|
| | | Rev: 5’TAATACGACTCACTATAGGGTATCCGCTCACAATTCCACA 3’ |
| 12 | DsrHPX12 | Fw: 5’ TAATACGACTCACTATAGGGTTCTGGTGTTTGCCATCGTA 3’<br>Rev: 5’ TAATACGACTCACTATAGGGCAGGATGTTCTGCTCGTTGA 3’ |
| 13 | SOD1 | Fw: 5' TGGGAGCACGCTTACTAT 3’<br>Rev: 5' GCTGTCGACTTTTGGCTA 3’ |
| 14 | SOD3 | Fw: 5' GATCGTCCGTCGCTATTA 3’<br>Rev: 5' CCTGCAAGCGTTATCTTC **3’** |
| 15 | Venom allergen | Fw: 5' GTTTTACGGAGTAACCAAGA 3’<br>Rev: 5' TACGGCATAGTTACAAACTG 3’ |
| 16 | Ovuline | Fw: 5'GAATGTAAGGGCATCATC3’<br>Rev: 5’AGATAGCGATTCTCTACGAA3’ |
| 17 | Sulfhydryl oxidase | Fw: 5' TTTTACAACTCGTACTGTGG 3’<br>Rev: 5' TCACTGGTTCTCCTATCTTC 3’ |
| 18 | Actin | Fw: 5’ TGCGTGACATCAAGGAGAAG 3’<br>Rev: 5’GATTCCATACCCAGGAACGA 3’ |

| S.No. | Gene Name | Primer Sequence |
|---|---|---|
| 1 | HPX2 | Fw: CACGAAGCTAAAAATTGTCC<br>Rev: AAGAGATGCTCCAGATCGTA |
| 2 | HPX3 | Fw: AGTTCTTCACGGTTCATCAC<br>Rev: GATCTTCTCCAGGTGTTTTG |
| 3 | HPX8 | Fw: AAGCTGTACCAAGAAGCTC<br>Rev: GCTCCTGATACTTCGTTAAA |
| 4 | HPX10 | Fw: AAGAAGGTTGACGGTACAGA<br>Rev: CAACGTTAGCAGTCGTATCA |
| 5 | HPX15 | Fw: CTATTGTCGCAACAGTACGA<br>Rev: CGGTAAACTGTCCATCATTT |
| 6 | Acp62f | Fw: 5' TACACGCTAGTGGTTTTGC 3'<br>Rev: : 5' AGCAAGATGTGGAAGGAATA 3' |
| 7 | HPX12 | Fw: GAACAGTGCCACCGATACCT<br>Rev: CCGAGATAATAGGGCAACCA |
| 8 | P-450 | Fw: TCATAATGTGGTGTACAAGC<br>Rev: ACCTAATCTCTATCGGTGTG |
| 9 | Catalase | Fw: 5' CCCAACTATTTCCCGAAC 3'<br>Rev: 5'CAGGTGTCCCACAATGTT3' |
| 10 | GST | Fw: 5' AACGGGTCGTCGATTACT 3'<br>Rev: 5' AGGTCGAACTGGAATGCT 3' |
| 11 | DsrLacz | Fw: 5' TAATACGACTCACTATAGGGGAGTCAGTGAGCGAGGAAG 3' |

| | | |
|---|---|---|
| | | Rev: 5'TAATACGACTCACTATAGGGTATCCGCTCACAATTCCACA 3' |
| 12 | DsrHPX12 | Fw: 5' TAATACGACTCACTATAGGGTTCTGGTGTTTGCCATCGTA 3'<br>Rev: 5' TAATACGACTCACTATAGGGCAGGATGTTCTGCTCGTTGA 3' |
| 13 | SOD1 | Fw: 5' TGGGAGCACGCTTACTAT 3'<br>Rev: 5' GCTGTCGACTTTTGGCTA 3' |
| 14 | SOD3 | Fw: 5' GATCGTCCGTCGCTATTA 3'<br>Rev: 5' CCTGCAAGCGTTATCTTC **3'** |
| 15 | Venom allergen | Fw: 5' GTTTTACGGAGTAACCAAGA 3'<br>Rev: 5' TACGGCATAGTTACAAACTG 3' |
| 16 | Ovuline | Fw: 5'GAATGTAAGGGCATCATC3'<br>Rev: 5'AGATAGCGATTCTCTACGAA3' |
| 17 | Sulfhydryl oxidase | Fw: 5' TTTTACAACTCGTACTGTGG 3'<br>Rev: 5' TCACTGGTTCTCCTATCTTC 3' |
| 18 | Actin | Fw: 5' TGCGTGACATCAAGGAGAAG 3'<br>Rev: 5'GATTCCATACCCAGGAACGA 3' |
